# CorridorKit: Open-source software for geometric comparison of skull base surgical corridors

**DOI:** 10.64898/2026.11.13.766552

**Authors:** Abhinav Bachu, Raghav Rajesh, Anand V. Germanwala

## Abstract

CorridorKit is open-source research software for comparing rigid surgical access corridors in physical image coordinates. Its Python engine represents finite instruments, circular portals, target samples, and protected geometry, and keeps evaluated paths linked to their reached samples, insertion depths, and clearance states through command-line, desktop, and 3D Slicer interfaces. In a reproducible 413-sample phantom, two portals reached 94 and 267 samples, with 326 in their union. Numerical verification included 82 analytical angular comparisons and 4,720 voxel-distance queries. In a public sellar CT example, conditional target coverage rose from 0 to 533 of 536 voxels as the modeled sphenoidotomy opening radius increased from 4 to 8 mm.

## 1. Motivation and significance

Comparing skull-base approaches requires specifying which instrument can reach which part of a target through a constrained opening. Instrument length, diameter, entry position, and surrounding anatomy all affect access. Linking a coverage result to its constituent paths makes these relationships available for inspection and repeatable experiments.

3D Slicer and SlicerIGT provide established infrastructure for medical-image review and intervention research [1,2]. Procedure-specific software supports laser-ablation planning, pedicle-screw placement, implant navigation, and stereoelectroencephalography [3–6]. Neurosurgical trajectory methods have addressed deep brain stimulation, keyhole insertion, and electrode placement [7–12]. Bériault et al. modeled cylindrical trajectories against multimodal segmentations [8], while Shamir et al. incorporated proximity-based risk and placement uncertainty [9].

Quantitative skull-base studies compare exposure and instrument maneuverability after anatomical preparation [13–17]. Volume of surgical freedom and volume of operative maneuverability extend this measurement framework into three dimensions [18,19]. MorphoNeuro provides an open-source Blender add-on for corridor morphometry [20]. Computer-based analysis of endonasal access also has a substantial history: de Notaris et al. used CT-based three-dimensional models to assess bony landmarks and bone removal [21]. The visuo-operative angle relates a surgical trajectory to the plane of target exposure in reconstructed skull-base corridors [22].

CorridorKit contributes a reusable implementation that combines finite instrument length and radius, conservative portal fit, retained trajectory witnesses, and set-based target coverage. These components let researchers connect a modeled constraint to the paths and target samples it affects. The software complements exposure and orientation measurements by making instrument-dependent reachability an explicit computational object. This article describes version 0.3.1 for technically supported surgeons and researchers.

## 2. Software description

### 2.1. Software architecture

The Python package separates typed domain models, image input/output, geometry, analysis, application services, anatomical adapters, visualization, and export (Figure 1). Pydantic contracts define cases and results. The command-line interface generates synthetic cases, runs analyses, and reports environment health. An optional Qt/VTK desktop application provides linked slices, three-dimensional inspection, configuration editing, asynchronous analysis, and export.

**Figure 1.**
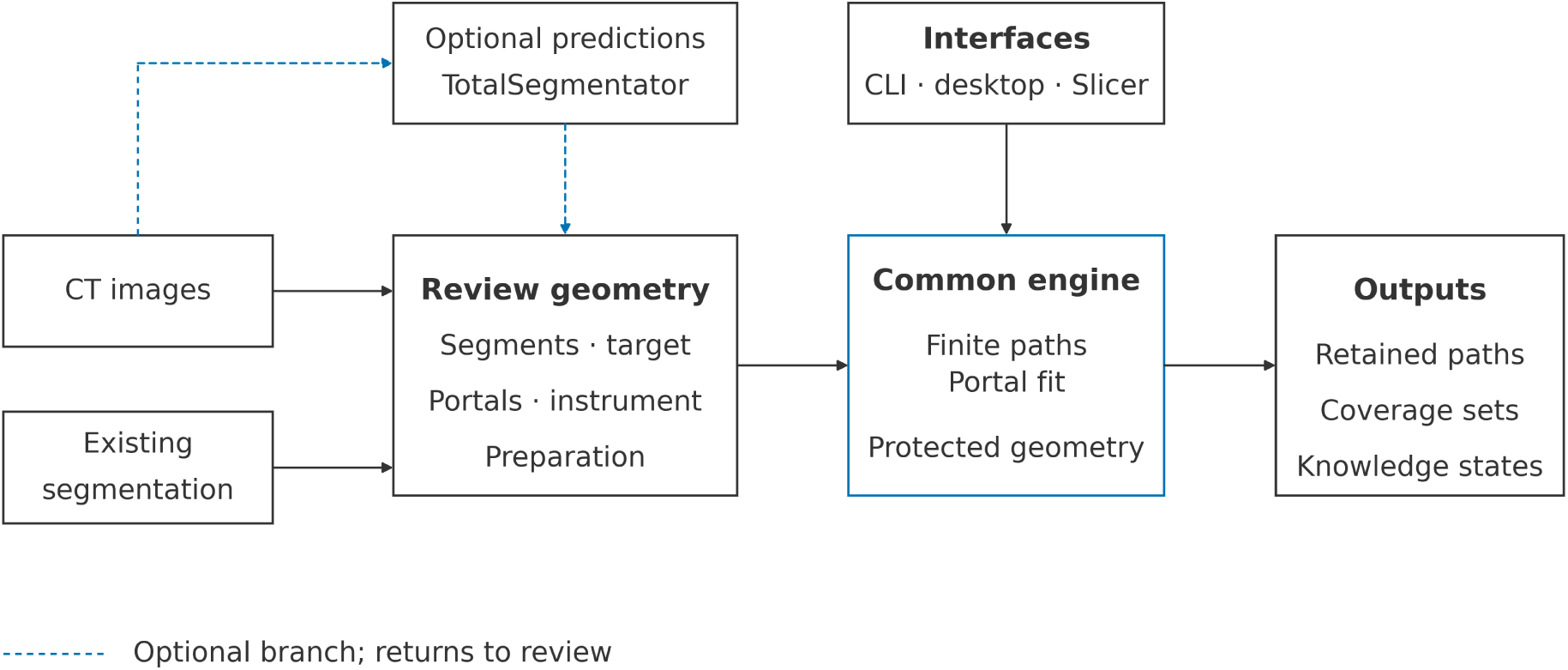
Software architecture and data flow. CT images and existing segmentation enter geometry review. Optional predictions return to review, while CLI, desktop, and Slicer interfaces share the common engine. Outputs include retained paths, coverage sets, and knowledge states.

Parameter changes invalidate displayed results until analysis is repeated. During development, the authors used Cursor (Anysphere) and ChatGPT (OpenAI) to assist with writing code; all code was directed, reviewed, tested, and verified by the authors, who take full responsibility for the software.

The scripted 3D Slicer module supports CT selection, markups, segmentation review, candidate trajectories, linked views, scene persistence, and reports. Its segment bridge exports the full CT grid, converts Slicer’s array ordering to engine ordering, and preserves the world-RAS affine. Targets exceeding the bridge’s 2,000-foreground-voxel limit are rejected explicitly. A managed external TotalSegmentator process can generate initial labels [23], using its nnU-Net foundation [24]. Users review the structures needed for their experiment before analysis; manually prepared segmentations enter through the same geometry contracts.

### 2.2. Software functionalities

#### 2.2.1. Physical coordinates and finite instruments

Geometry is expressed in RAS millimeters, denoting right, anterior, and superior axes. Affines map voxel indices into physical space. Image loaders normalize declared coordinate conventions and units and check frame compatibility. Registration and resampling are explicit operations. Targets comprise physical points or mask-derived voxel centers. Full voxel representations may carry cell-volume weights from the absolute affine determinant; sparse targets are reported as sample counts.

For entry e, unit direction d, and target sample x, axial insertion is t = (x − e)·d and perpendicular distance is ‖(x − e) − td‖. A reach candidate requires nonnegative insertion within instrument length and perpendicular distance within working-tip radius plus target tolerance. Shaft radius is separate from target tolerance. The inserted instrument is a finite capsule ending at the insertion depth required for that target. Collision and aperture checks conservatively use the larger of shaft and tip radius (Figure 2).

**Figure 2.**
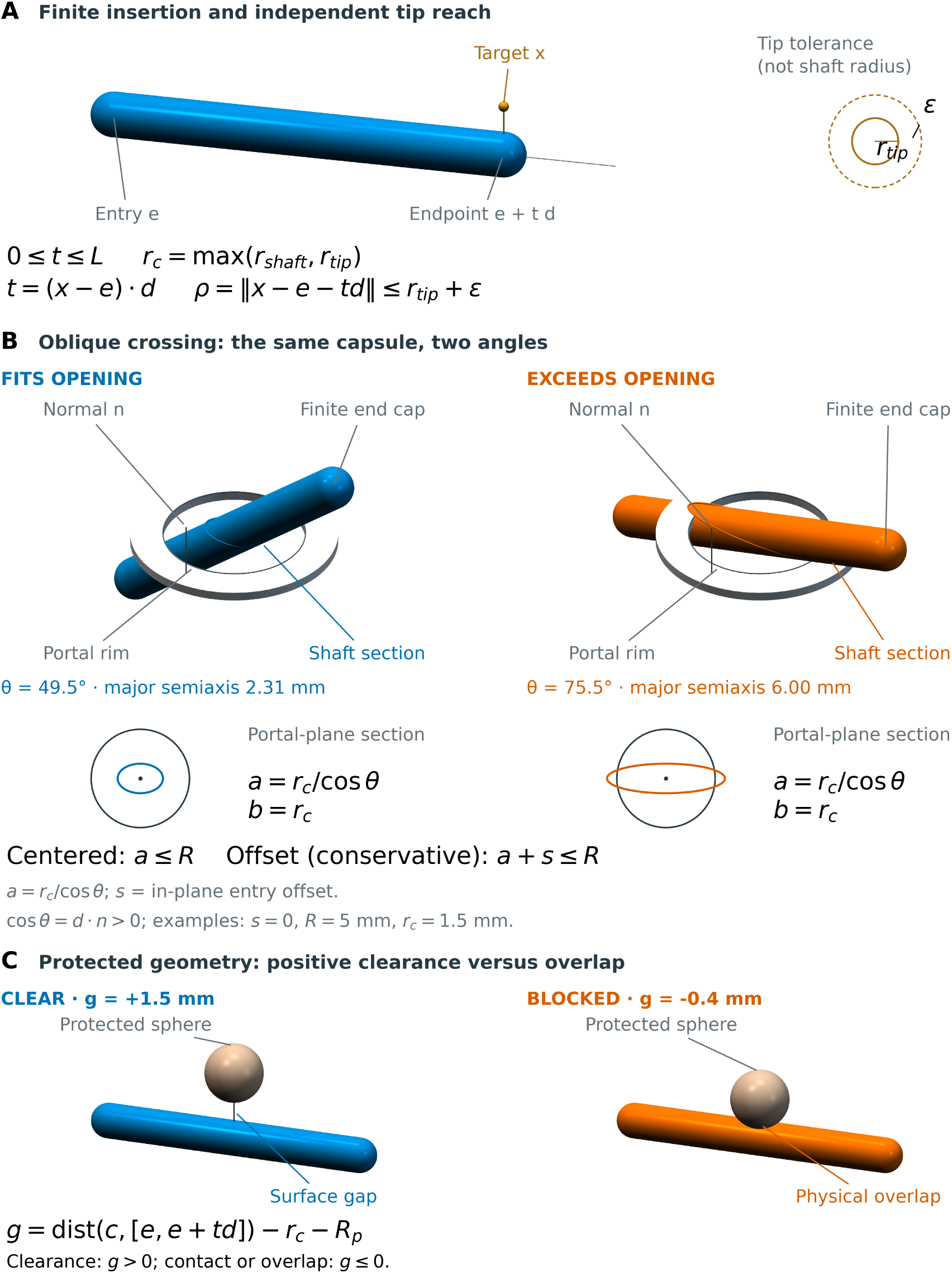
Finite-instrument geometry. (A) A finite insertion ends at e + t d; the inset shows tip reach and target tolerance. (B) Two crossing angles produce elliptical shaft sections that fit or exceed a circular portal. The major semiaxis is a = r_c/cos(θ), R is the portal radius, and s is the in-plane entry offset; centered fit requires a ≤ R and the conservative offset condition is a + s ≤ R. (C) A protected sphere has positive surface clearance g > 0 in the clear example and overlaps the capsule in the blocked example.

A portal is a circular disk with a forward normal. The engine checks forward traversal and conservative fit of the oblique instrument footprint. Protected structures can be analytical spheres or affine voxel masks. Sphere clearance is computed directly; masks represent closed physical cells. Conservative voxel bounds can be refined near a boundary using a finite candidate-cell budget. Budget exhaustion retains conservative blockage. Required geometry outside a mask’s field of view invokes the configured policy. Supplementary Methods S1–S3 specify these procedures.

#### 2.2.2. Paths and coverage

The engine evaluates a configured angular grid with optional target-directed and budgeted adaptive directions. Each trajectory retains its direction, reached target indices, insertion depths, clearance information, and rejection or abstention reason. A retained feasible trajectory is a witness to the configured result. The interface distinguishes sampled-unreached targets from targets whose assessment is unavailable.

Approach-specific target-index sets support union, intersection, and difference without double counting. These operations are exact on the evaluated sets. Simultaneous-access analysis additionally checks angular separation and finite capsule-to-capsule clearance. Missing, unknown, unsupported, or out-of-field protected anatomy invokes explicit knowledge states and the selected analysis policy. Unsupported meshes remain visible as unsupported inputs.

#### 2.2.3. Measurements and export

The reported comparisons use retained paths and target coverage for configurations containing protected geometry. Solid-angle verification is confined to supported unobstructed center-entry configurations, using geometry-aware interval-union integration. This keeps each reported measurement tied to its tested numerical procedure.

Exports record the normalized configuration, software and schema versions, input identifiers, paths, coverage sets, and analysis states. Images, predictions, scenes, and reports remain on the workstation unless exported. A skull-base neurosurgeon co-author (A.V.G.) reviewed the corridor model, research workflow, and interpretation of geometric results.

## 3. Illustrative examples

### 3.1. Reproducible workflows

From the versioned source directory, a minimal workflow is:

~~~
python -m pip install -e.
corridorkit synthetic case.json
corridorkit analyze case.json result.json
corridorkit doctor
~~~

The desktop interface is available through the optional desktop dependencies. Its planning phantom is a separate fixture from the minimal command-line synthetic case. For reviewed-segment analysis in Slicer, users load CT and aligned segmentations, identify target and entry geometry, select protected structures, and configure instrument and portal dimensions. Approach-specific bone-removal masks record the modeled preparation. Returned target categories are approach-only, shared, sampled-unreached, or unavailable.

For model-assisted analysis, Slicer exports CT to a private workspace and starts the external segmentation worker. Predictions are mapped to the CT grid for review. Model weights are retrieved separately on first use. Researchers can vary instrument dimensions, portal size, or preparation and rerun the same case to compare the resulting sets and witnesses.

### 3.2. Synthetic coverage example

The planning phantom contains a CT-like volume, a 413-sample ellipsoidal target, two portals, and three protected spheres. Portal A and Portal B carry nominal endoscopic endonasal and transmaxillary labels. The display shell supplies visual context; collision geometry is defined by the protected spheres.

Portal A yielded 27 feasible trajectories reaching 94 samples; Portal B yielded 77 reaching 267. Their partition comprised 59 A-only, 232 B-only, 35 shared, and 87 sampled-unreached samples, with zero unavailable samples (Table 2). The union reached 326 distinct samples. The desktop workflow retained 104 selectable witnesses and cleared stale paths after configuration changes.

**Table 1.**
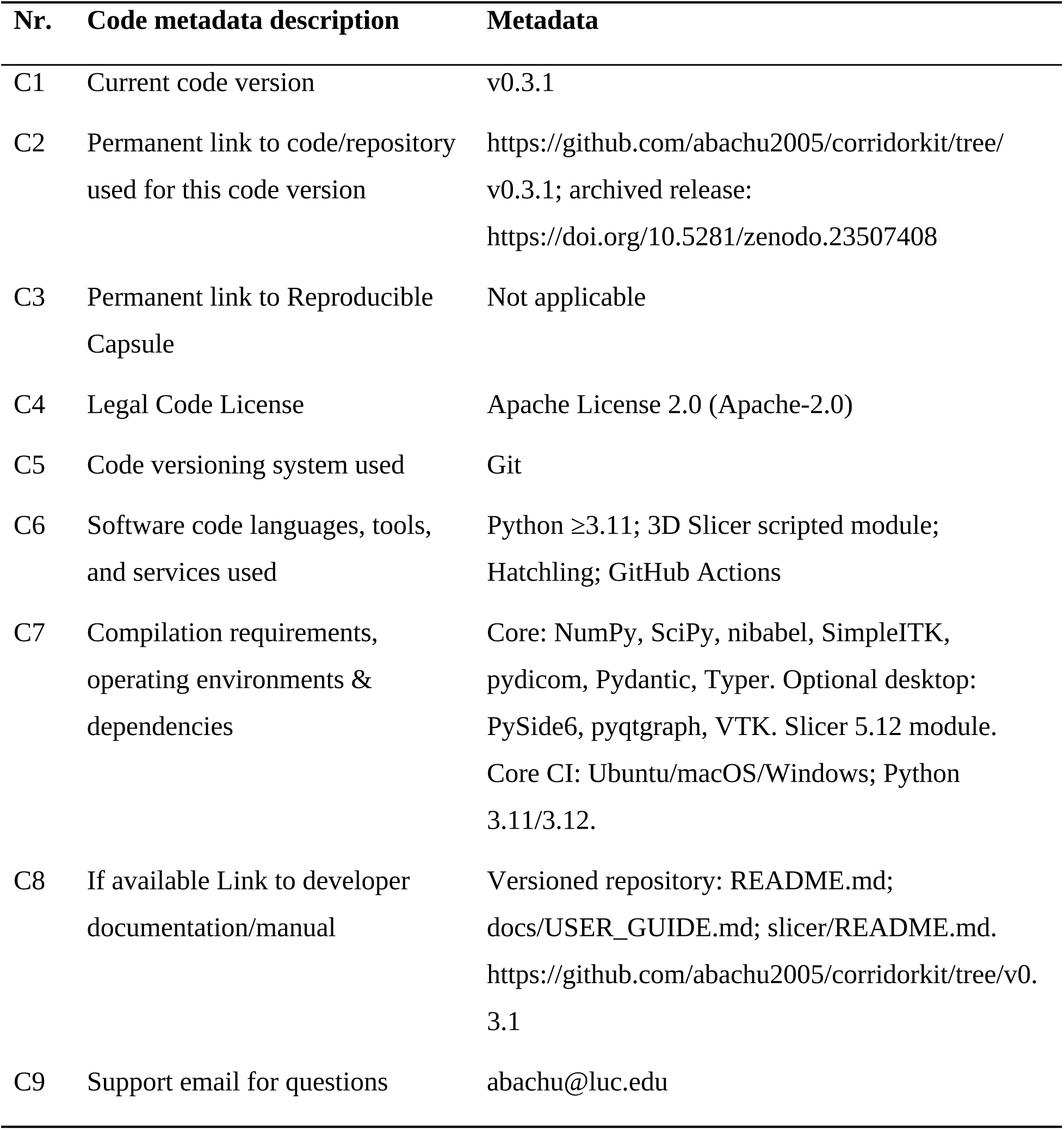
Code metadata.

**Table 2.** Synthetic planning phantom with 413 target samples.

| Measure | Count | Interpretation |
|---|---|---|
| Portal A feasible trajectories | 27 | Evaluated finite paths |
| Portal B feasible trajectories | 77 | Evaluated finite paths |
| Portal A reached samples | 94 | 59 A-only + 35 shared |
| Portal B reached samples | 267 | 232 B-only + 35 shared |
| A-only samples | 59 | Partition category |
| B-only samples | 232 | Partition category |
| Shared samples | 35 | Partition category |
| Sampled-unreached samples | 87 | Partition category |
| Unavailable samples | 0 | Partition category |
| Alternative union | 326 | Distinct samples reached by either portal |

The partition is 59 A-only + 232 B-only + 35 shared + 87 sampled-unreached + 0 unavailable = 413. Alternative union = 326 distinct reached samples.

### 3.3. Illustrative sellar CT example

We used public HaN-Seg case_01 [25,26] to examine how modeled bone preparation changes endonasal access. Figure 3 shows CT and segmentation inspection and bilateral portal configuration in the desktop workspace; Figures 4 and 5 show the patient-derived anatomy and modeled preparation. The complete 536-voxel pituitary mask (333 mm3) supplied the target. Six expert masks represented the optic chiasm, both optic nerves, both carotid artery contours, and brainstem; bone was thresholded from the CT at 300 HU or above. We preserved the native grid and derived bilateral nasal-vestibule portals from enclosed CT air. A 180-mm instrument with 2-mm shaft radius was evaluated along 289 trajectories per portal.

**Figure 3.**
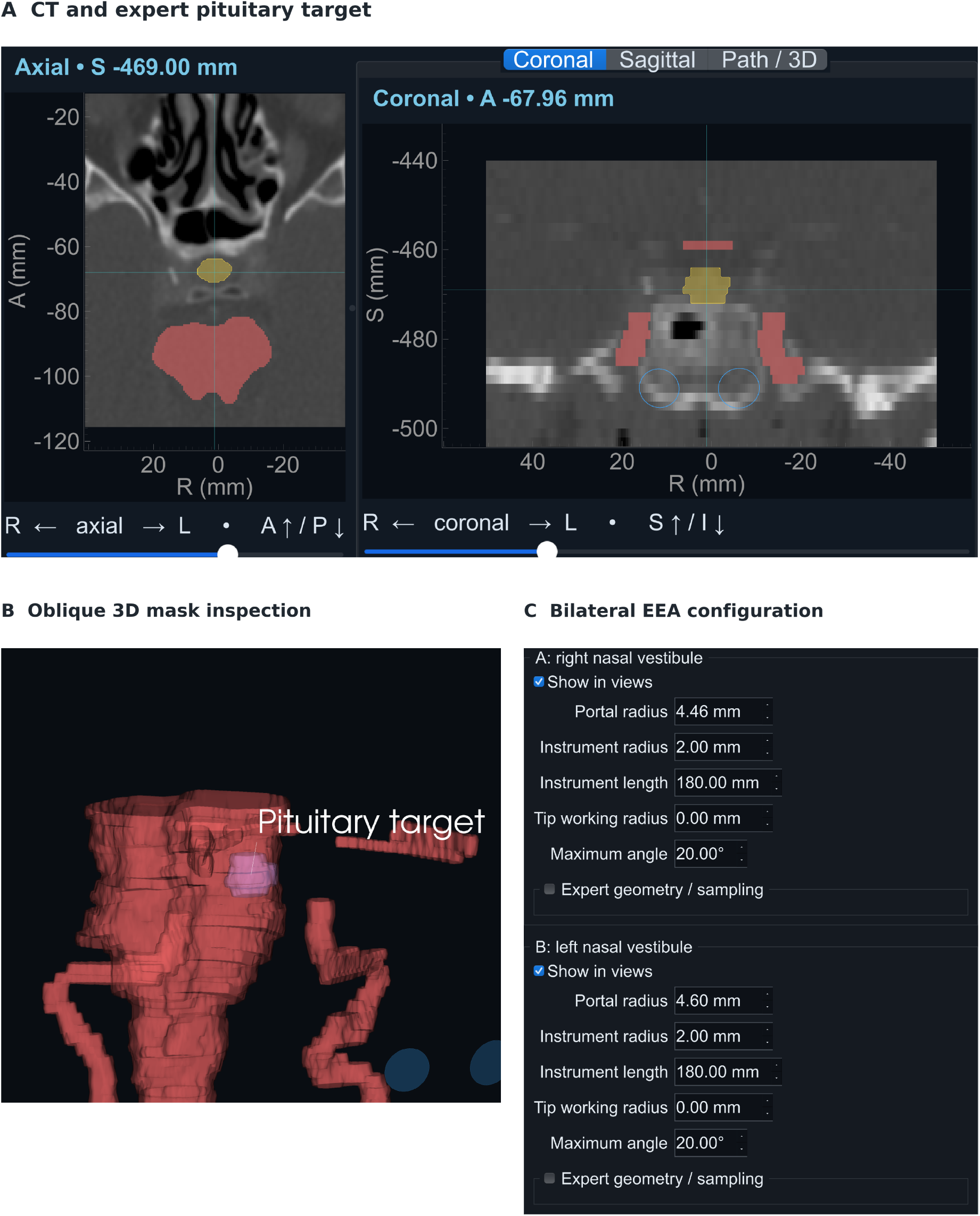
CorridorKit workspace on public HaN-Seg case_01. (A) Linked axial and coronal CT with the expert pituitary target (536 native voxels) and protected segmentations. (B) Oblique 3D inspection of the pituitary (purple), carotid arteries, optic nerves, optic chiasm, and brainstem. (C) Bilateral illustrative EEA nasal-vestibule aperture and instrument configuration. Incomplete anatomy limits the stored results to conditional scope; conditional witnesses remain unselectable and are shown in Figure 5. Source: HaN-Seg [25,26].

**Figure 4.**
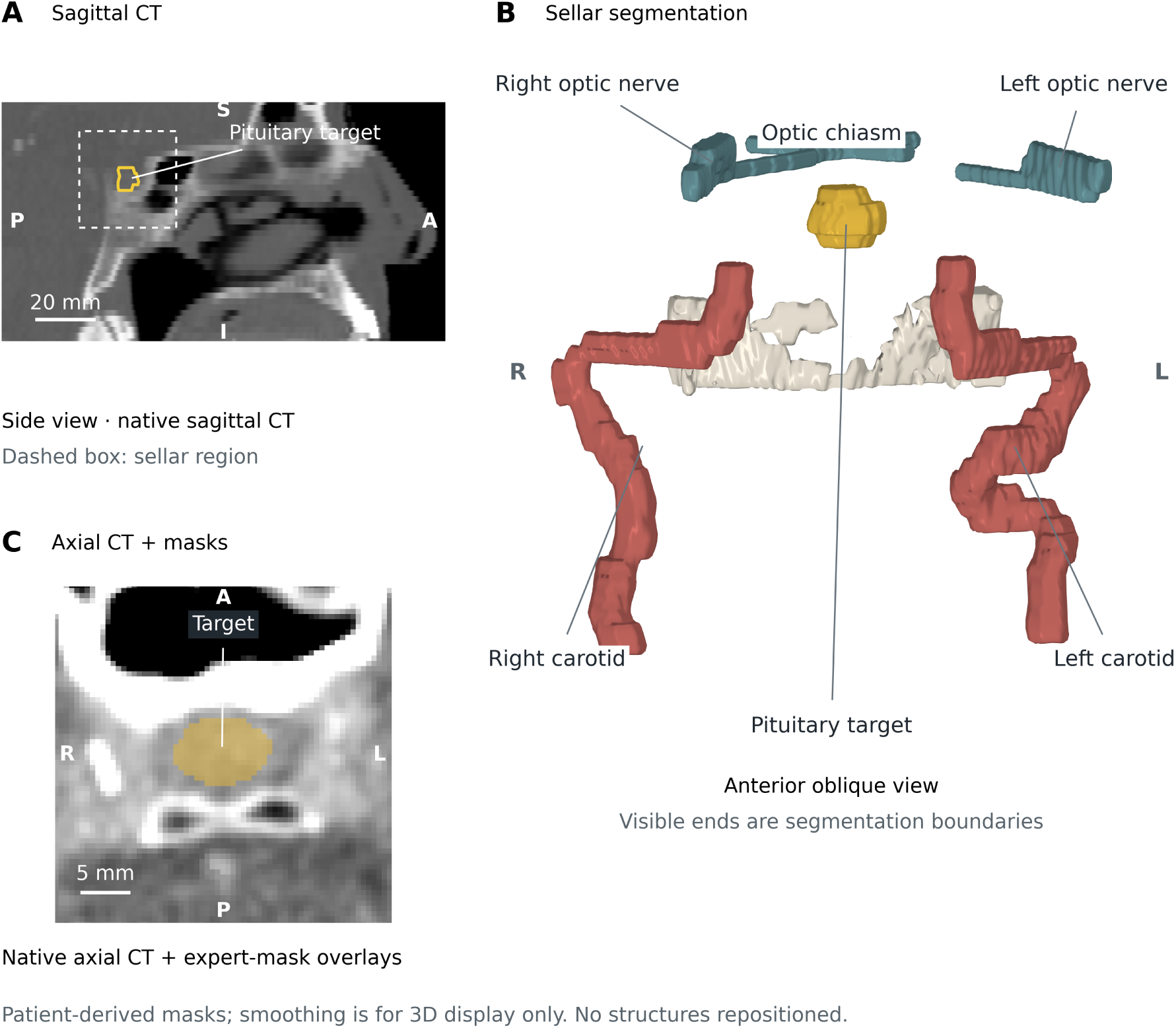
Patient-derived pituitary and protected anatomy in HaN-Seg case_01. (A) Native sagittal CT locates the target and boxed sellar region. (B) Anterior-oblique expert-mask surfaces with posterior bone context; visible ends follow segmentation boundaries, not image cropping. Surface smoothing and the existing tiny-component carotid filter affect display only; retained masks stay in native positions. (C) Native axial CT with pituitary and carotid mask overlays. Source: HaN-Seg [25,26].

**Figure 5.**
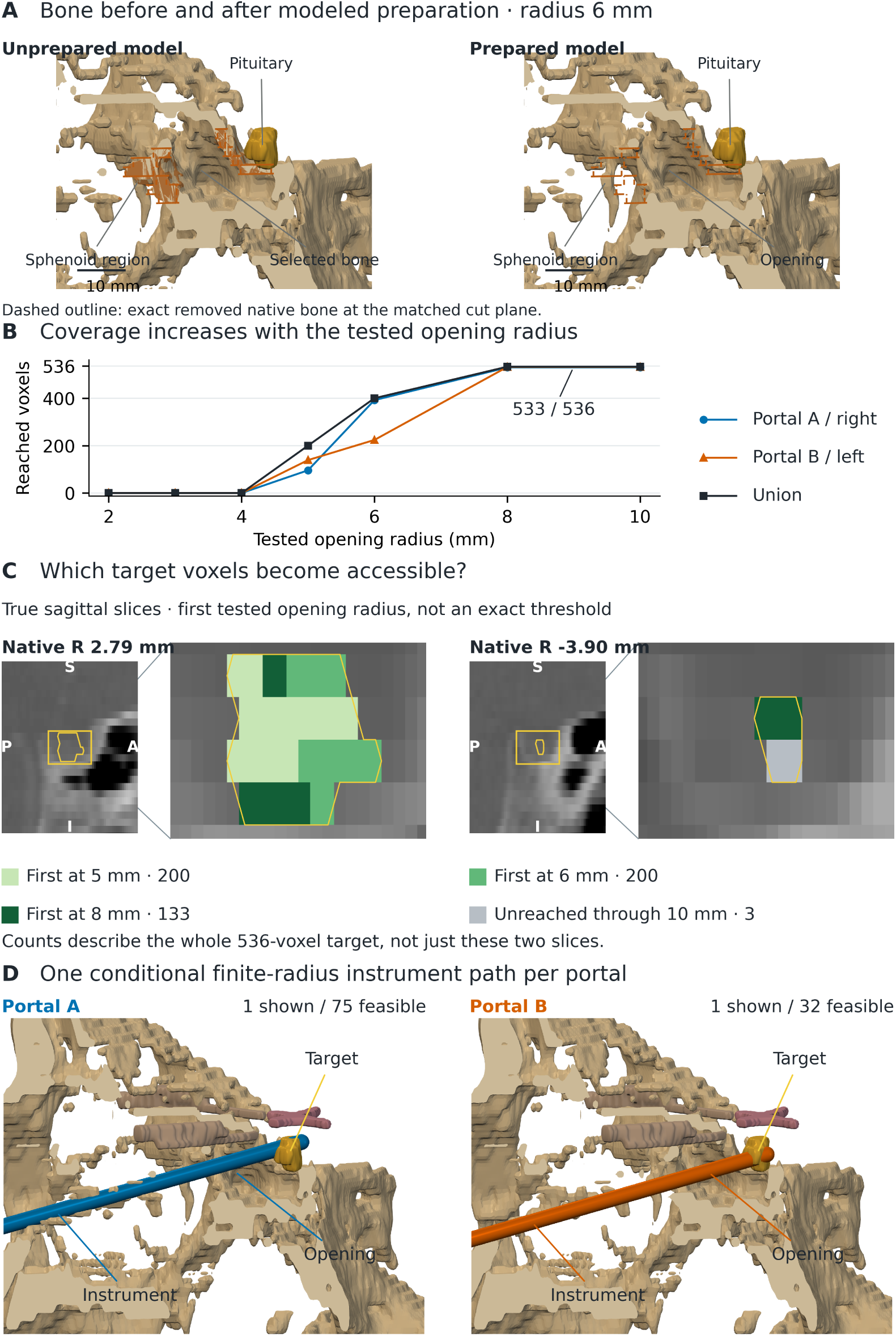
Modeled preparation and conditional target access in HaN-Seg case_01. (A) Matched CT-derived cutaways before and after modeled Portal A preparation at radius 6 mm. The identical dashed outlines mark the exact removed native bone-cell boundary at the R=3.5 mm display cut plane; orange marks selected bone before removal and gold marks the expert pituitary target. Ivory cut faces and expanded anterior CT bone provide display context. (B) Stored conditional sampled-trajectory coverage; the union reaches 533 of 536 target voxels at tested radii 8 to 10 mm. Portal A is right and Portal B is left. (C) Native sagittal target cells colored by first tested opening radius; counts describe the whole 536-voxel target. (D) One stored conditional finite-radius instrument witness per portal with optic apparatus and prepared bone context. Witnesses maximize portal-exclusive reached target IDs, then total reached IDs, breaking ties by the smallest zero-based stored trajectory index. Surface smoothing is display-only, not clearance-measurement geometry. Source: HaN-Seg [25,26].

With bone undeclared, 530 right-portal and 532 left-portal voxels were reached, a union of 533. Adding CT-derived bone reduced this union to zero. When every thresholded voxel was represented as rigid, target-reaching trajectories contacted bone beginning in the nasal cavity. In the skull-base model, nasal septal and turbinate bone anterior to the preparation plane was declared unknown and displaceable, while the sphenoid face and sellar floor remained protected. This unprepared configuration also reached zero target voxels (Table 3).

**Table 3.**
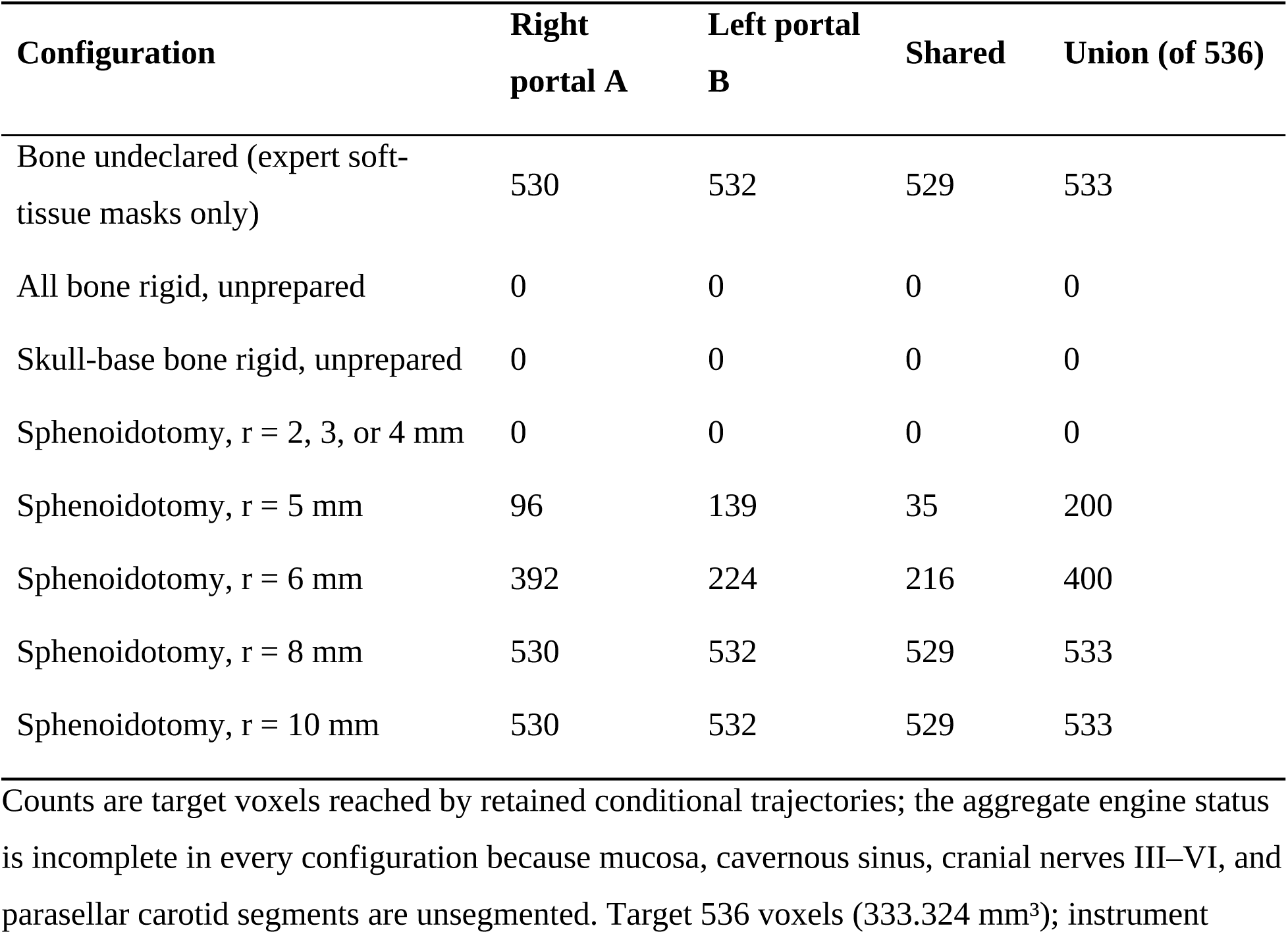
Sellar target access by bone model and modeled preparation in HaN-Seg case_01.

| **Configuration** | **Right portal A** | **Left portal B** | **Shared** | **Union (of 536)** |
|---|---|---|---|---|
| Bone undeclared (expert soft-tissue masks only) | 530 | 532 | 529 | 533 |
| All bone rigid, unprepared | 0 | 0 | 0 | 0 |
| Skull-base bone rigid, unprepared | 0 | 0 | 0 | 0 |
| Sphenoidotomy, r = 2, 3, or 4 mm | 0 | 0 | 0 | 0 |
| Sphenoidotomy, r = 5 mm | 96 | 139 | 35 | 200 |
| Sphenoidotomy, r = 6 mm | 392 | 224 | 216 | 400 |
| Sphenoidotomy, r = 8 mm | 530 | 532 | 529 | 533 |
| Sphenoidotomy, r = 10 mm | 530 | 532 | 529 | 533 |

Counts are target voxels reached by retained conditional trajectories; the aggregate engine status is incomplete in every configuration because mucosa, cavernous sinus, cranial nerves III–VI, and parasellar carotid segments are unsegmented. Target 536 voxels (333.324 mm<sup>3</sup>); instrument

A modeled sphenoidotomy, an approach-specific cylinder of bone removal about each portal axis from the sphenoid face through the sellar floor, restored access as the opening widened. Opening radii of 2–4 mm reached no voxels. A 5-mm opening radius reached 200 voxels (96 right, 139 left), 6 mm reached 400 (392 right, 224 left), and 8 and 10 mm reached 533, matching the bone-undeclared result. The 4–8-mm opening radii correspond to 8–16-mm diameters, ranging from approximately the median transverse sellar opening reported after transsphenoidal surgery (8.8 mm) [27] to a wider exposure. At intermediate openings the portals were complementary: 61 and 176 voxels were reached only from the right portal at 5 and 6 mm, and 104 and 8 only from the left. Native target slices show the first tested opening radius reaching each voxel (Figure 5C), and representative retained paths show finite-instrument access at 6 mm (Figure 5D). An independent finite-segment calculation checked all 3,330 reached trajectory–voxel pairs at 6 mm against remaining bone and expert masks with zero disagreements. The minimum conservative clearance lower bound was 0.140 mm, and replay preserved all discrete path classifications and reached sets. Supplementary Method S4.2, Table S5, and Figure S2 give the construction and per-configuration counts.

### 3.4. Numerical and engineering verification

We tested unobstructed angular integration against 82 analytical comparisons covering spherical caps, rotations, finite-length bands, clipped lenses, and target unions. All met the benchmark’s 1% relative-error criterion; the maximum observed error was 0.0001204123%.

Voxel refinement was first checked using 530 synthetic queries and 120 public queries from cases P001/P002/P004. We then applied the same bounded protocol to all 118 geometry-eligible NasalSeg v2 image/label pairs [28,29], with 40 queries per pair (4,720 total). Twelve of the 130 pairs failed image–label physical-geometry agreement. Refinement recovered 2,164 clear classifications from conservative blockage, each checked against a candidate-cell distance reference. In 1,190 budget-exhausted queries, the blocked fallback was retained. The checks recorded zero false-clear classifications, clearance-bound violations, field-of-view-policy changes, or fallback changes. Supplementary Tables S1–S3 provide the input accounting, numerical results, and angular stress test.

Release verification for v0.3.1 comprised 657 local test passes with no skips, source-distribution and clean-environment wheel checks, and desktop-dependency tests on macOS. Eight continuous-integration jobs passed on the release commit, including Python 3.11/3.12 core testing on Ubuntu, macOS, and Windows. Native desktop interaction and Slicer 5.12.3 integration on macOS were verified for the desktop and Slicer modules, which v0.3.1 leaves unchanged; native Slicer checks covered landmark invalidation, rotated anisotropic coordinate conversion, five-category overlays, external-engine execution, asynchronous analysis, and scene-reference round-trip (Supplementary Table S4).

## 4. Impact

CorridorKit enables controlled experiments on instrument-dependent access. Researchers can hold the target and protected geometry fixed while varying instrument length, diameter, portal radius, entry position, or modeled bone removal. Retained paths show where a parameter change adds or removes access, and coverage partitions quantify the affected target samples. These experiments can guide the choice of configurations for anatomical studies and support comparisons across implementations.

The phantom illustrates the value of retaining these sets. Although Portal B reached more samples overall, Portal A added 59 samples beyond the Portal B set. The union captured this incremental contribution while counting the 35 shared samples once. The same operations can compare preparation stages, instrument profiles, or alternative entry geometries on a common target grid. In the sellar example, the instrument and nasal portals were held fixed while sphenoid-opening radius changed. The sweep quantified the target coverage added by preparation and showed that the portal with greater coverage changed with opening size.

The image interfaces make these experiments reusable in existing research workflows. Investigators can inspect path witnesses beside CT and reviewed segmentations, save the scene, and export the configuration for replay. The separate engine also supports parameter sweeps and comparison with tracked instrument paths or anatomical exposure measurements [13–22]. Its typed physical-space inputs provide a basis for adapting the same finite-access calculations to other rigid-instrument research problems.

The model represents rigid instruments and static anatomy. Anatomical interpretation depends on segmentation, registration, target and portal definitions, and represented preparation. In the sellar example, CT-threshold bone at 2-mm slice thickness under-resolves thin septal and turbinate bone, nasal bone anterior to the sphenoid face is treated as displaceable, and the sphenoidotomy is a rule-defined cylinder rather than a surgeon-drawn resection. Its results remain conditional on unsegmented mucosa, cavernous sinus, cranial nerves, and parasellar carotid segments. Finite sampling leaves continuous reachability unresolved. The voxel-refinement reference shares broad-phase candidate selection with the implementation. Protected-geometry grid quadrature remains exploratory and is excluded from the reported comparisons.

## 5. Conclusions

CorridorKit combines finite rigid-instrument analysis, aperture constraints, retained trajectory witnesses, and set-based coverage in an open, reusable software package. Its image interfaces and reproducible examples enable researchers to study how declared instrument and corridor geometry shape target access.

## Code and data availability

CorridorKit v0.3.1 is available under Apache-2.0 at https://github.com/abachu2005/corridorkit/tree/v0.3.1 and https://doi.org/10.5281/zenodo.23507408. Dependencies retain their licenses. HaN-Seg and NasalSeg v2 are available separately under their stated dataset licenses [26,29]. The sellar-example scripts are in the repository at examples/sellar_hanseg (https://github.com/abachu2005/corridorkit/tree/v0.3.1/examples/sellar_hanseg). Verification protocols and summary results are tracked in the repository (docs/VALIDATION.md, docs/evidence/ and research/); the NasalSeg v2 eligibility audit, including the 118 geometry-eligible image/label pairs, is research/nasalseg-v2-audit.json. Per-query benchmark outputs are not publicly archived.

## Ethics statement

This work used synthetic and publicly available deidentified data, with no recruitment, private-record access, chart review, or identity linkage.

## CRediT authorship contribution statement

Abhinav Bachu: Conceptualization, Methodology, Software, Validation, Investigation, Data curation, Visualization, Writing – original draft, Writing – review & editing. Raghav Rajesh: Investigation, Visualization, Writing – original draft, Writing – review & editing. Anand V. Germanwala: Methodology, Validation, Supervision, Writing – review & editing.

## Declaration of generative AI and AI-assisted technologies in the manuscript preparation process

During the preparation of this work, the authors used Cursor (Anysphere), an AI-assisted coding tool, and ChatGPT (OpenAI) to assist with language editing and with writing software and analysis code. AI tools did not independently generate content; all text, code, and analyses were directed, reviewed, edited, and verified by the authors, who take full responsibility for the content of the publication.

## References

[1] Fedorov A, Beichel R, Kalpathy-Cramer J, Finet J, Fillion-Robin JC, Pujol S, et al. 3D Slicer as an image computing platform for the Quantitative Imaging Network. Magn Reson Imaging 2012;30(9):1323–41. 10.1016/j.mri.2012.05.001.

[2] Ungi T, Lasso A, Fichtinger G. Open-source platforms for navigated image-guided interventions. Med Image Anal 2016;33:181–6. 10.1016/j.media.2016.06.011.

[3] Yeniaras E, Fuentes DT, Fahrenholtz SJ, Weinberg JS, Maier F, Hazle JD, et al. Design and initial evaluation of a treatment planning software system for MRI-guided laser ablation in the brain. Int J Comput Assist Radiol Surg 2014;9(4):659–67. 10.1007/s11548-013-0948-x.

[4] Muralidharan V, Swaminathan G, Devadhas D, Joseph BV. Patient-specific interactive software module for virtual preoperative planning and visualization of pedicle screw entry point and trajectories in spine surgery. Neurol India 2018;66(6):1766–70. 10.4103/0028-3886.246281.

[5] Chen X, Xu L, Wang H, Wang F, Wang Q, Kikinis R. Development of a surgical navigation system based on 3D Slicer for intraoperative implant placement surgery. Med Eng Phys 2017;41:81–9. 10.1016/j.medengphy.2017.01.005.

[6] Narizzano M, Arnulfo G, Ricci S, Toselli B, Tisdall M, Canessa A, et al. SEEG assistant: a 3DSlicer extension to support epilepsy surgery. BMC Bioinformatics 2017;18(1):124. 10.1186/s12859-017-1545-8.

[7] Brunenberg EJL, Vilanova A, Visser-Vandewalle V, Temel Y, Ackermans L, Platel B, et al. Automatic trajectory planning for deep brain stimulation: a feasibility study. Med Image Comput Comput Assist Interv 2007;10(Pt 1):584–92. 10.1007/978-3-540-75757-3_71.

[8] Bériault S, Al Subaie F, Collins DL, Sadikot AF, Pike GB. A multi-modal approach to computer-assisted deep brain stimulation trajectory planning. Int J Comput Assist Radiol Surg 2012;7(5):687–704. 10.1007/s11548-012-0768-4.

[9] Shamir RR, Joskowicz L, Tamir I, Dabool E, Pertman L, Ben-Ami A, et al. Reduced risk trajectory planning in image-guided keyhole neurosurgery. Med Phys 2012;39(5):2885–95. 10.1118/1.4704643.

[10] Trope M, Shamir RR, Joskowicz L, Medress Z, Rosenthal G, Mayer A, et al. The role of automatic computer-aided surgical trajectory planning in improving the expected safety of stereotactic neurosurgery. Int J Comput Assist Radiol Surg 2015;10(7):1127–40. 10.1007/s11548-014-1126-5.

[11] Sparks R, Vakharia V, Rodionov R, Vos SB, Diehl B, Wehner T, et al. Anatomy-driven multiple trajectory planning (ADMTP) of intracranial electrodes for epilepsy surgery. Int J Comput Assist Radiol Surg 2017;12(8):1245–55. 10.1007/s11548-017-1628-z.

[12] Wankhede A, Madiraju L, Siampli E, Fischer E, Cleary K, Oluigbo C, et al. Validation of a novel path planner for stereotactic neurosurgical interventions-A retrospective clinical study. Int J Med Robot 2022;18(6):e2458. 10.1002/rcs.2458.

[13] Roth J, Singh A, Nyquist G, Fraser JF, Bernardo A, Anand VK, et al. Three-dimensional and 2-dimensional endoscopic exposure of midline cranial base targets using expanded endonasal and transcranial approaches. Neurosurgery 2009;65(6):1116–28; discussion 1128– 30. 10.1227/01.NEU.0000360340.85186.7A.

[14] Wilson DA, Williamson RW, Preul MC, Little AS. Comparative analysis of surgical freedom and angle of attack of two minimal-access endoscopic transmaxillary approaches to the anterolateral skull base. World Neurosurg 2014;82(3-4):e487–93. 10.1016/j.wneu.2013.02.003.

[15] Elhadi AM, Almefty KK, Mendes GA, Kalani MYS, Nakaji P, Dru A, et al. Comparison of surgical freedom and area of exposure in three endoscopic transmaxillary approaches to the anterolateral cranial base. J Neurol Surg B Skull Base 2014;75(5):346–53. 10.1055/s-0034-1372467.

[16] Lin BJ, Ju DT, Hsu TH, Chen YA, Chung TT, Liu WH, et al. Quantitative comparison of endoscopically assisted endonasal, sublabial and transorbital transmaxillary approaches to the anterolateral skull base. Clin Otolaryngol 2021;46(1):123–30. 10.1111/coa.13559.

[17] Agosti E, Saraceno G, Rampinelli V, Raffetti E, Veiceschi P, Buffoli B, et al. Quantitative Anatomic Comparison of Endoscopic Transnasal and Microsurgical Transcranial Approaches to the Anterior Cranial Fossa. Oper Neurosurg 2022;23(4):e256–66. 10.1227/ons.0000000000000312.

[18] Houlihan LM, Naughton D, Preul MC. Volume of Surgical Freedom: The Most Applicable Anatomical Measurement for Surgical Assessment and 3-Dimensional Modeling. Front Bioeng Biotechnol 2021;9:628797. 10.3389/fbioe.2021.628797.

[19] Tariciotti L, Rodas A, Patel B, Zohdy YM, Revuelta Barbero JM, De Andrade EJ, et al. Volume of Operative Maneuverability as a New Measurement in Neuroanatomical Research: A Methodological Quantitative Study and Translational Use in the Operating Room. World Neurosurg 2025;196:123761. 10.1016/j.wneu.2025.123761.

[20] Sinosi FA, Rodriguez Rubio R. An Open Source Tool to Conduct 3-Dimensional Morphometric Analysis of Microneurosurgical Corridors: Technical Note and Anatomical Validation Study Using Surface Scanning Techniques. World Neurosurg 2025;202:124378. 10.1016/j.wneu.2025.124378.

[21] de Notaris M, Solari D, Cavallo LM, Enseñat J, Alobid I, Soria G, et al. The use of a three-dimensional novel computer-based model for analysis of the endonasal endoscopic approach to the midline skull base. World Neurosurg 2011;75(1):106–13. 10.1016/j.wneu.2010.09.033.

[22] Tariciotti L, Rodas A, Zohdy YM, Revuelta Barbero JM, De Andrade EJ, Patel B, et al. The visuo-operative angle: computational 3D assessment of surgical trajectories toward irregular target areas in skull base corridors. J Neurosurg 2026;144(6):1463–76. 10.3171/2025.9.JNS251518.

[23] Wasserthal J, Breit HC, Meyer MT, Pradella M, Hinck D, Sauter AW, et al. TotalSegmentator: Robust Segmentation of 104 Anatomic Structures in CT Images. Radiol Artif Intell 2023;5(5):e230024. 10.1148/ryai.230024.

[24] Isensee F, Jaeger PF, Kohl SAA, Petersen J, Maier-Hein KH. nnU-Net: a self-configuring method for deep learning-based biomedical image segmentation. Nat Methods 2021;18(2):203–11. 10.1038/s41592-020-01008-z.

[25] Podobnik G, Strojan P, Peterlin P, Ibragimov B, Vrtovec T. HaN-Seg: The head and neck organ-at-risk CT and MR segmentation dataset. Med Phys 2023;50(3):1917–27. 10.1002/mp.16197.

[26] [dataset] Podobnik G, Strojan P, Peterlin P, Ibragimov B, Vrtovec T. HaN-Seg: The head and neck organ-at-risk CT & MR segmentation dataset, Zenodo, Version 1.0; 2023. 10.5281/zenodo.7442914.

[27] Yahia-Cherif M, Delpierre I, Hassid S, De Witte O. Bony Regeneration of the Sella after Transsphenoidal Pituitary Surgery. World Neurosurg 2016;88:497–502. 10.1016/j.wneu.2015.10.073.

[28] Zhang Y, Wang J, Pan T, Jiang Q, Ge J, Guo X, et al. NasalSeg: A Dataset for Automatic Segmentation of Nasal Cavity and Paranasal Sinuses from 3D CT Images. Sci Data 2024;11(1):1329. 10.1038/s41597-024-04176-1.

[29] [dataset] Zhang Y, Wang J, Pan T, Jiang Q, Ge J, Guo X, et al. NasalSeg Dataset for Nasal Cavity and Paranasal Sinuses Segmentation from CT Images, Zenodo, Version 2; 2024. 10.5281/zenodo.13893419.

