## Supplementary Material for "CorridorKit: Open-source software for geometric comparison of skull base surgical corridors"

*Contents: S1–S5, Tables S1–S5, and Figures S1–S2. Bracketed reference numbers follow the reference list of the main article; the two references cited here are listed at the end.*

### S1. Computational inputs

We evaluated target samples, trajectories, angular comparisons, and voxel-distance queries. NasalSeg v2 contains 130 public CT scans and five sinonasal labels covering the bilateral maxillary sinuses, bilateral nasal cavities, and nasopharynx (Zhang et al., Scientific Data 2024;11:1329; doi:10.1038/s41597-024-04176-1). The dataset is available at https://doi.org/10.5281/zenodo.13893419 under CC BY 4.0.

We required exact-ID image/label pairing, finite three-dimensional arrays, integral labels within 0–5, nonsingular matching RAS affines (absolute tolerance 0.0001; relative tolerance zero), and orthogonal voxel axes. Of 130 pairs, 118 met these computational requirements and 12 failed physical-geometry agreement (Table S1).

Excluded identifiers were P025, P039, P042, P074, P089, P096, P100, P101, P105, P108, P118, and P129. The initial development subset comprised P001/P002/P004. The subsequent 118-case run included these cases. We group repeated queries by source identifier and report computational counts.

### S2. Finite instruments and target coverage

All engine coordinates are in RAS millimeters. Image affines map voxel indices to physical positions. Full voxel-center targets may carry cell weights $|\det(A_{3}\times_{3})|$ in $mm^3$; sparse targets are reported as sample counts. Coordinate normalization occurs during import, with registration and resampling handled explicitly.

For entry e, unit direction d, and target x, axial insertion and perpendicular distance are:

$$t = (x - e) \cdot d$$

$$\rho = \|(x - e) - td\|$$

A candidate requires $t \geq 0$, $t \leq L$, and $\rho \leq r_{tip} + \varepsilon_{target}$. L denotes instrument length, $r_{tip}$ the working-tip radius, and $\varepsilon_{target}$ target tolerance. Shaft radius is separate from target tolerance. Collision and aperture checks use $r_{collision} = \max(r_{shaft}, r_{tip})$. The finite capsule extends from e to e + td for that target. Protected spheres are checked analytically.

The portal is a physical circular disk with a forward normal and radius. The engine checks forward traversal and conservative fit of the oblique footprint. For reached target-index sets A and B, unique coverage is $|A \cup B|$, shared coverage is $|A \cap B|$, and incremental coverage is $|A \setminus B|$ or $|B \setminus A|$. Simultaneous analysis additionally checks angular separation and finite capsule clearance.

The direction search combines an angular grid with optional target-directed and budgeted adaptive witnesses. Results retain the path direction, reached indices, insertion depths, and analysis state. Unknown, unsupported, or out-of-field protected anatomy invokes the selected knowledge policy. This distinguishes unavailable assessment from sampled-unreached targets.

### S3. Numerical verification

#### S3.1. Angular integration

We tested geometry-aware interval-union integration for supported unobstructed center-entry configurations with at most 32 target points. The benchmark used seed 20261001 and a 1% relative-error acceptance criterion. Integration defaults were relative tolerance $10^{-5}$, absolute tolerance $10^{-12}$ sr, at most 100,000 evaluations, and a 5-second budget.

The 82 comparisons comprised 16 original comparisons, 48 randomized caps, eight frame rotations, four finite-length annuli, three clipped lenses, and three target unions. References used spherical-cap formulas, rotation invariance, finite axial bands, spherical-lens geometry, and inclusion–exclusion. Relative error was |estimate − reference|/|reference|. All 82 met the criterion, with maximum observed error 0.0001204123%.

Version 0.3.1 also implements spherical-cap grid quadrature with Voronoi area weights for protected-structure configurations. Target-directed and adaptive witnesses receive no quadrature weight. This routine emits a grid estimate without an automatic convergence gate; it is excluded from the reported protected-geometry comparisons. Table S3 gives the separate off-axis 11° grid stress test, whose finest grid had 11.368% relative error. Supported unobstructed integration and this exploratory grid routine are distinct procedures.

#### S3.2. Voxel clearance refinement

The coarse voxel backend uses conservative physical-space bounds and interpolation guards. Near-boundary refinement checks closed affine cells under a default budget of 128 candidate cells. Reported refined positive clearance is capped at 0.1 mm. Budget exhaustion retains conservative blockage, and refinement preserves the configured field-of-view policy.

The reference uses bounded convex least squares for segment-to-cell distance and support-function lower bounds, with numerical tolerance $10^{-8}$ mm. A clearance-bound violation occurs when reported positive clearance exceeds the optimizer's upper distance bound after instrument-radius subtraction, subject to that comparison tolerance. The reference and implementation share broad-phase candidate selection.

The initial run included 530 synthetic and 120 public queries. Per public case, two anchors × four targets × two radii (0 and 1 mm) produced 16 segment queries; 12 near-boundary centers × two radii (0 and 0.25 mm) produced 24 point-capsule queries. The 118-case run therefore contained 1,888 anchor–target and 2,832 near-boundary queries (4,720 total). All 2,164 recovered-clear public classifications were reference checked. Table S2 reports the results. The synthetic suite included 30 out-of-field queries; the public suite included zero.

### S4. Reproducible examples

#### S4.1. Planning phantom

The planning phantom contains a 413-sample ellipsoidal target on a 2-mm grid, two 4-mm-radius portals, and three protected spheres. Both instruments have length 100 mm and shaft radius 0.8 mm, with target tolerance 1 mm. CT-like shell and airspaces provide display context. Portal A and Portal B have nominal endonasal and transmaxillary labels.

Sampling uses a 35° maximum angle, two polar steps, eight azimuth steps, target-directed sampling with at most 96 target witnesses, no adaptive levels, and no portal-offset rings. The source constructor is src/corridorkit/synthetic/planning.py. Call write_planning_phantom, read the returned case, and call analyze_case using its parent as base_directory. The phantom differs from the minimal command-line synthetic fixture.

#### S4.2. Illustrative sellar CT example

We used case_01 from the 42-case public training subset of HaN-Seg, whose reference masks were curated from expert manual annotations (Podobnik et al., Med Phys. 2023;50:1917–1927; doi:10.1002/mp.16197). The source CT grid is 1024 × 1024 × 202 with spacing 0.557617 × 0.557617 × 2.0 mm. Spatial units were interpreted as millimeters because the NRRD headers omit a unit field. CT and masks were cropped without resampling to a 181 × 289 × 43 grid covering the nasal vestibules, sella, and represented protected structures. The crop retained every target voxel and the full optic and brainstem masks.

The target comprises all 536 pituitary voxel centers, with volume 333.324 mm³ from the affine determinant. Protected voxel masks represent the optic chiasm, bilateral optic nerves, bilateral carotid artery contours, and brainstem. We declared three further anatomy categories unknown: nasal mucosa, septal cartilage, and turbinate soft tissue; cavernous sinus and cranial nerves III–VI; and cavernous/supraclinoid carotid segments above the dataset contours. The exploratory setting allow_unknown_anatomy = true retains conditional per-trajectory outputs while the aggregate engine status remains incomplete; all coverage counts are unions of target indices from feasible trajectories flagged conditional.

The portal rule selects the most inferior axial slice between S = −497 and −481 mm containing enclosed air below −400 HU on both sides of the target-centroid sagittal plane, within A = 15–38 mm. Each portal is centered on the largest inscribed circle in its enclosed-air region (Figure S1). The selected slice is S = −491 mm. Right and left centers are RAS (11.710,19.583,−491.000) and (−6.134,19.583,−491.000) mm, with radii 4.461 and 4.598 mm. Portal normals point toward the target centroid.

All configurations use instrument length 180 mm, shaft radius 2 mm, zero additional working-tip radius, and target tolerance 1 mm. Sampling uses a 20° cone, five polar steps, 24 azimuth steps, at most 64 target-directed witnesses, one adaptive level with a 128-direction refinement budget, and center-only portal entry. The engine evaluated 289 trajectories per portal.

For the bone-undeclared baseline, for each of the 7,556 reached trajectory–voxel pairs, we independently computed the minimum distance from the finite insertion segment to every represented protected voxel center. Subtracting half the voxel diagonal and the instrument radius gives a conservative clearance lower bound for the closed cells. The calculation found zero negative-clearance disagreements.

Bone was thresholded at ≥300 HU from the unmodified CT crop (228,734 voxels). In the all-bone model, every thresholded voxel is rigid protected geometry. In the skull-base model, voxels anterior to the coronal plane A = −36.16 mm, 5 mm anterior to the more anterior sphenoid-face point, are excluded (145,972 voxels retained); this septal and turbinate bone is declared unknown as displaceable tissue, not free space.

On each portal-to-target-centroid axis, sampled at 0.25 mm, the sphenoid face is the start of the penultimate ≥300-HU run and the sellar floor is the last run: insertion depths 63.25 and 83.00 mm (right) and 68.25 and 81.25 mm (left). The modeled sphenoidotomy removes skull-base bone within radius r of the axis, from max(1 mm, r) anterior to the face, enclosing an anterior wall inclined up to 45°, to the deepest target voxel along the axis (94.64 and 94.39 mm). Each removal is a subset of the bone mask and is attached to its approach as a virtual bone removal; its reviewed flag records author inspection of the rule output, not clinician approval. We evaluated r = 2, 3, 4, 5, 6, 8, and 10 mm with the instrument fixed (Table S5).

Coverage first appeared at a 5-mm opening with the fixed 2-mm shaft radius and conservative voxel enclosure of 1.07 mm (half the voxel diagonal). For the 6-mm configuration, we independently computed exact distances from every finite insertion segment to all remaining bone and expert-mask voxel centers. We subtracted the half-diagonal enclosure and instrument radius to obtain conservative clearance lower bounds for all 3,330 reached trajectory–voxel pairs. All were positive; the minimum was 0.1403 mm. Replaying the stored unprepared and 6-mm cases preserved all discrete outputs and reached sets, with floating fields agreeing within an absolute tolerance of $10^{-10}$. Figure S2 shows the insertion depths of bone contact in the unprepared models.

To reproduce the example, install CorridorKit with nibabel, SimpleITK, SciPy, and matplotlib and run the scripts in examples/sellar_hanseg (https://github.com/abachu2005/corridorkit/tree/v0.3.1/examples/sellar_hanseg). fetch_members.py retrieves the case_01 CT and seven masks from https://doi.org/10.5281/zenodo.7442914 by byte range; prepare_case.py builds the native-grid case and the bone-undeclared baseline; run_bone_preparation.py evaluates both bone models and the opening sweep; and verify_bone_preparation.py repeats the checks above. The directory README lists the commands. The verifier script (verify_bone_preparation.py) includes the exact finite-segment clearance check. Verification protocols and summary results are tracked in the repository (docs/VALIDATION.md, docs/evidence/ and research/); per-query benchmark outputs are not publicly archived.

### S5. Software verification and availability

Release verification for v0.3.1 comprised 14 local groups totaling 657 test passes with no skips, source-distribution and clean-environment wheel checks, and desktop-dependency tests on macOS; native desktop interaction and Slicer integration on macOS were verified for modules that v0.3.1 leaves unchanged. Eight continuous-integration jobs passed on the release commit, including Python 3.11/3.12 core checks on Ubuntu, macOS, and Windows. Table S4 lists the checks.

The source is https://github.com/abachu2005/corridorkit/tree/v0.3.1, archived at https://doi.org/10.5281/zenodo.23507408 under Apache-2.0. The source distribution includes code, tests, documentation, and deterministic fixtures. Optional desktop and Slicer dependencies support native interface reproduction. Verification protocols and summary results are tracked in the repository (docs/VALIDATION.md, docs/evidence/ and research/); the NasalSeg v2 eligibility audit, including the 118 geometry-eligible image/label pairs, is research/nasalseg-v2-audit.json. Per-query benchmark outputs are not publicly archived. The sellar-example scripts are in examples/sellar_hanseg. HaN-Seg source data are available at https://doi.org/10.5281/zenodo.7442914 under CC BY-NC-ND 4.0; NasalSeg v2 and optional model weights are retrieved separately.

**Table S1.** Computational input accounting

| Stage | Count | Meaning |
|---|---|---|
| Public image/label pairs | 130 | NasalSeg v2 source identifiers |
| Excluded pairs | 12 | Physical image–label geometry mismatch |
| Engineering-eligible pairs | 118 | All included in full numerical query replay |
| Initial development subset | 3 | P001/P002/P004; 40 queries each |
| Full query replay | 4,720 | 118 × 40; includes development cases |

Excluded identifiers are listed in S1. The full run includes the three development cases.

**Table S2.** Voxel refinement outcomes

| Quantity | Synthetic | Initial public | Full public |
|---|---|---|---|
| Total queries | 530 | 120 | 4,720 |
| Recovered-clear queries | 185 | 62 | 2,164 |
| Reference/oracle queries | 495 | 62 | 2,164 |
| Budget-exhausted queries | 5 | 30 | 1,190 |
| False-clear classifications | 0 | 0 | 0 |
| Clearance-bound violations | 0 | 0 | 0 |
| FOV-policy changes | 0 | 0 | 0 |
| Budget-fallback changes | 0 | 0 | 0 |
| Out-of-field queries | 30 | 0 | 0 |

Initial public: 3 cases × 40 queries. Full public: 118 cases × 40 queries, including the initial cases. The repeated synthetic suite is counted once. FOV = field of view.

**Table S3.** Off-axis grid quadrature stress test

| Grid | Directions | Estimated sr | Relative error % |
|---|---|---|---|
| 5 × 24 | 97 | 0.268377429 | 111.4105 |
| 9 × 48 | 385 | 0.191698163 | 51.0075 |
| 17 × 96 | 1,537 | 0.158150985 | 24.5812 |
| 33 × 192 | 6,145 | 0.141377396 | 11.3680 |

Reference: 0.12694612240263908 sr. Grid dimensions are polar steps × azimuth steps; duplicate pole directions are consolidated. These results characterize the exploratory grid routine; reported protected-geometry comparisons use paths and coverage sets.

**Table S4.** Software verification for v0.3.1

| Check | Result | Scope |
|---|---|---|
| Local test groups | 657 passed; 0 skipped | Numerical group and 13 isolated groups |
| Source distributions | Passed | Source-only contents; source unchanged during build |
| Installed wheel | Passed | Clean-environment install and CLI smoke test |
| Slicer archive | Passed | Package integrity; module unchanged in v0.3.1 |
| Native desktop | Passed | 104 selectable paths; stale-result invalidation |
| Native Slicer | Passed | Slicer 5.12.3 on macOS |
| Landmark invalidation | Passed | Native Slicer integration |
| Rotated anisotropic coordinates | Passed | KJI-to-IJK/RAS export |
| Coverage categories | Passed | Five-category synthetic overlay |
| External engine and dialog | Passed | Process execution and asynchronous workflow |
| Scene-reference round-trip | Passed | Save/reopen integration |
| Cross-platform CI | 8 jobs passed | Release commit; Python 3.11/3.12 on three OS families |

Native desktop and Slicer checks used macOS and cover modules unchanged in v0.3.1. Core continuous integration covered Python 3.11/3.12 on Ubuntu, macOS, and Windows.

**Table S5.** Sellar configurations: trajectories and removed bone

| Configuration | Feasible A / B | Bone-blocked A / B | Removed voxels A / B | Union |
|---|---|---|---|---|
| All bone rigid, unprepared | 0 / 0 | 154 / 153 | 0 / 0 | 0 |
| Skull-base bone rigid, unprepared | 0 / 0 | 154 / 153 | 0 / 0 | 0 |
| Sphenoidotomy, r = 2 mm | 0 / 0 | 154 / 153 | 154 / 219 | 0 |
| Sphenoidotomy, r = 3 mm | 0 / 0 | 154 / 153 | 329 / 459 | 0 |
| Sphenoidotomy, r = 4 mm | 0 / 0 | 154 / 153 | 700 / 752 | 0 |

| Configuration | Feasible A / B | Bone-blocked A / B | Removed voxels A / B | Union |
|---|---|---|---|---|
| Sphenoidotomy, r = 5 mm | 7 / 16 | 147 / 137 | 1,139 / 1,163 | 200 |
| Sphenoidotomy, r = 6 mm | 75 / 32 | 79 / 121 | 1,697 / 1,693 | 400 |
| Sphenoidotomy, r = 8 mm | 147 / 138 | 7 / 15 | 2,814 / 2,801 | 533 |
| Sphenoidotomy, r = 10 mm | 153 / 144 | 1 / 9 | 4,044 / 3,936 | 533 |

289 trajectories per portal; 135 (right) and 136 (left) miss the target geometrically in every configuration, and all other unreached trajectories are structure-blocked. Removed voxels are skull-base bone voxels inside each approach's removal cylinder. Union counts are target voxels of 536.

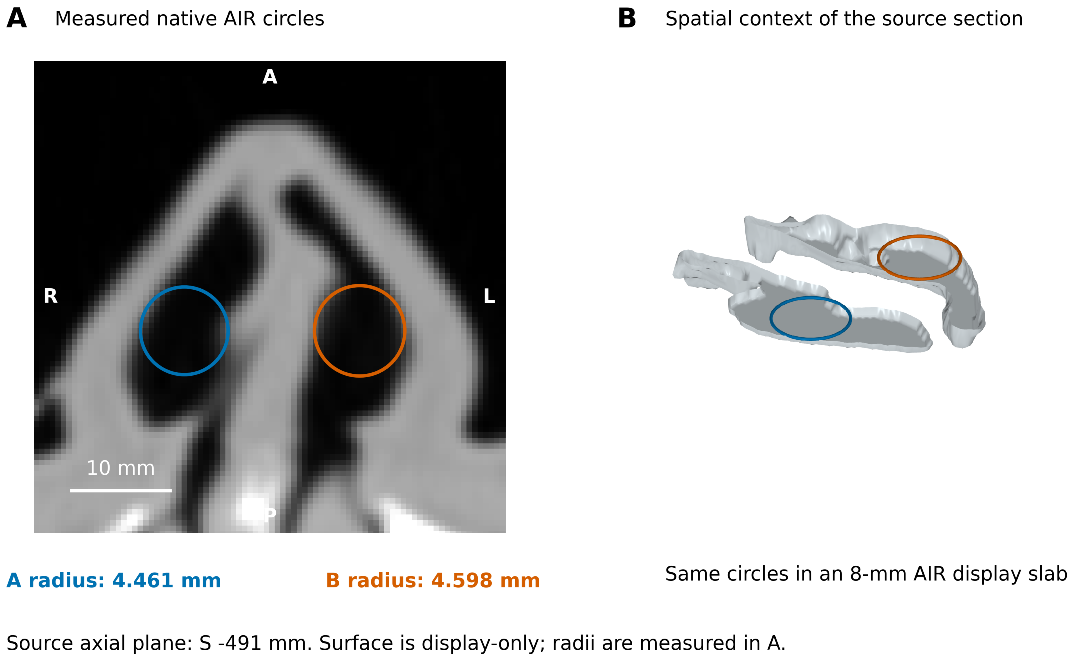


**Figure S1.** Portal-circle derivation. (A) Exact inscribed circles in native nasal AIR on the source axial plane S -491 mm: A 4.461 mm and B 4.598 mm. (B) The same axial circles in a display-only 8-mm AIR slab, with exterior-connected air excluded. These measured axial circles are not the tilted engine portal disks. Source: HaN-Seg [25,26].

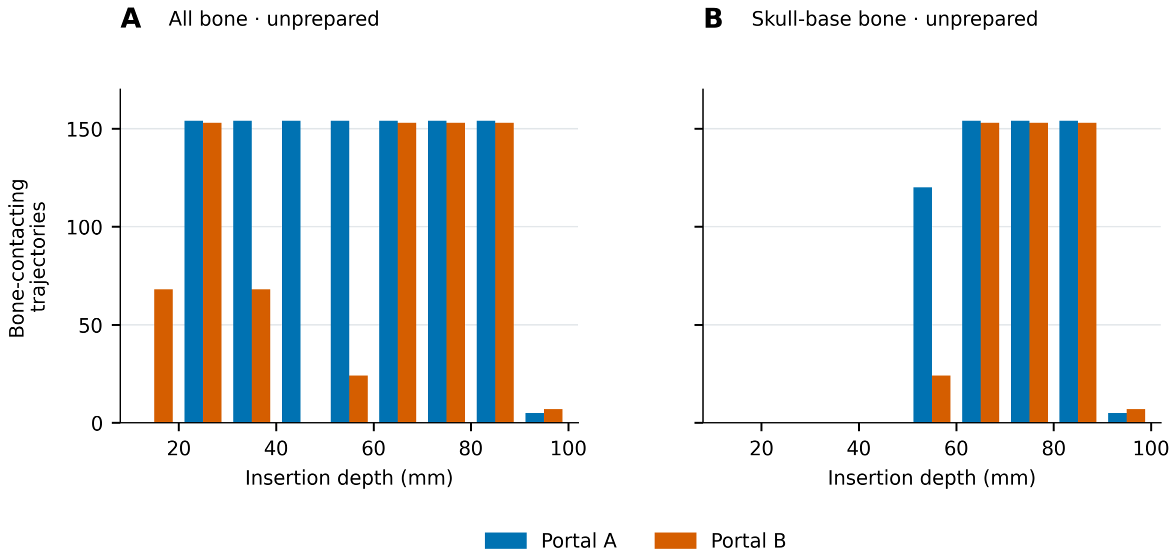


**Figure S2.** Stored unprepared bone-contacting trajectory counts in aligned insertion-depth bins for (A) all bone and (B) skull-base bone. Portal A and B retain consistent colors. Insertion depth uses 10-mm bins; an individual trajectory may contribute to multiple bins. Bins report contacting sampled trajectories, not independent continuous anatomical thresholds. Source: HaN-Seg [25,26].

### References

[25] Podobnik G, Strojan P, Peterlin P, Ibragimov B, Vrtovec T. HaN-Seg: The head and neck organ-at-risk CT and MR segmentation dataset. Med Phys 2023;50(3):1917–27. https://doi.org/10.1002/mp.16197.

[26] [dataset] Podobnik G, Strojan P, Peterlin P, Ibragimov B, Vrtovec T. HaN-Seg: The head and neck organ-at-risk CT & MR segmentation dataset, Zenodo, Version 1.0; 2023. https://doi.org/10.5281/zenodo.7442914.
